# Isoform-level discovery, quantification and fusion analysis from single-cell and spatial long-read RNA-seq data with Bambu-Clump

**DOI:** 10.1101/2024.12.30.630828

**Authors:** Andre Sim, Min Hao Ling, Ying Chen, Han Lu, Yi Xiang See, Arnaud Perrin, Ong Bee Leng Agnes, Elaine Yiqun Cao, Burton Chia, Jinyue Liu, Torsten Wüstefeld, Jay W. Shin, Jonathan Göke

## Abstract

Single cell and spatial transcriptomics have dramatically changed how we can profile RNA from heterogenous biological samples. Combining single cell and spatial profiling with long read RNA-Seq promises to enable the discovery and quantification of individual RNA isoforms at the single-cell level. However, highly multiplexed data such as from a single cell experiment only generates a limited number of reads for each cell, constituting a major challenge for transcript discovery and quantification with existing approaches that usually have limited power for samples with low sequencing depth.

Here we present Bambu-Clump, a computational method that performs transcript discovery and quantification from single cell and spatial long read RNA-Seq data using information from both each cell and the cell cluster. Using this approach, Bambu-Clump provides the most accurate transcript discovery compared to other existing methods, and improves transcript quantification compared to methods that rely on estimates derived from single cells. We apply Bambu-Clump to identify fusion transcripts in single-cells, compare 5’ and 3’ selection protocols, and identify novel isoform cell-type markers in spatial mouse brain data. Together, Bambu-Clump provides an easy-to-use, efficient, and accurate method for analysing individual isoform expression for single cells and cell clusters across multiple datasets and replicates from long read RNA-Seq.

## Introduction

Single-cell and spatial RNA-seq technology enables the profiling of the transcriptome for individual cells, cell types, and anatomic regions. Many samples used in RNA-seq are not uniform in nature, rather they are a heterogeneous melange of differing cell types and tissues, making them particularly relevant for these technologies ^1^. These include, but are not limited to tumours, early embryos and brain tissue, which all often have complex differentiation distinguishing cells from one another ^2–5^.

Due to alternative splicing, alternative promoters, or alternative polyadenylation, most genes can generate alternative isoforms, often with distinct expression patterns or function. Doing so results in transcriptome diversity between tissues and cells, and has been observed in medically relevant cases such as in cancer tissues and the brain ^6–8^. However, short-read single cell or spatial RNA sequencing produces reads derived from the 5’ or 3’ ends of RNA transcripts and like Smart-seq based methods are too short to discriminate between most alternative isoforms ^9^. As it is not possible to unambiguously determine which isoforms are expressed from these read fragments, the analysis of single cell RNA-Seq data is typically limited to quantifying the total RNA expression that is observed for each gene (“gene expression”) ^10^. To overcome this limitation, single cell and spatial transcriptome profiling can be combined with PacBio or Oxford Nanopore long-read RNA sequencing. As no RNA fragmentation is required, complete RNA molecules can be sequenced, enabling transcriptome analysis at isoform resolution. However, as in bulk long-read data ^11^, despite containing full-length reads, the presence of truncated reads creates ambiguity requiring the use of statistical methods to accurately identify and quantify novel transcripts.

Several methods have been developed that specifically perform transcript discovery and quantification for bulk long read RNA-Seq data ^8,12–14^. However, these methods are optimised for deep transcriptome profiling of small sample numbers, whereas single cell RNA-seq data generates low depth transcriptome profiling of thousands of cells from multiple samples. To address this, single cell long read RNA-Seq pipelines such as wf-single-cell or FLAMES have been developed that handle the alignment, barcode detection and demultiplexing, and then use existing methods for transcript discovery and quantification ^15–20^. In addition, dedicated single cell long read RNA-Seq methods have been developed for transcript discovery and quantification both at the single cell ^12,19,20^ and pseudo bulk level ^21^. However, these methods can be resource intensive, unable to account for sample and technology differences when applying to different technologies, not compatible with both single-cell and spatial data, and not able to track full length read counts.

In this work we present Bambu-Clump, a computational method that enables efficient analysis of isoform expression using long read single cell and spatial RNA-Seq data. Bambu-Clump supports the simultaneous analysis of multiple samples, it enables adjustable transcript discovery at the bulk and pseudo bulk level, it can be used for quantification of fusion transcripts, and it returns full length read support for all transcripts to support cell-marker validation. Bambu-Clump is implemented as part of the Bambu package ^13^ and integrated into a nextflow pipeline (Bambu-Pipe) directly from raw reads, with an optional mode for fusion isoform discovery in conjunction with JAFFAL ^22^. We evaluate and compare Bambu-Pipe on 10x single cell and spatial long read RNA-Seq data from human cancer cell lines, and mouse brain. We then apply Bambu-Pipe to discover fusion transcripts in single cells, compare isoform expression and full length read counts for 5’ and 3’ single cell RNA-Seq transcriptomic protocols, and to identify marker isoforms in spatial mouse brain data. Together, we provide an efficient, accurate, adjustable, and easy to use method for analysing transcript expression data for single cell and spatial long read RNA-Seq.

## Results

### Isoform-level analysis of single-cell and spatial long read RNA-Seq data with Bambu-clump and Bambu-pipe

To address the need for efficient analysis of highly multiplexed samples, such as single-cell, single-nuclei, or spatial data, we developed Bambu-Clump. Implemented as a module in the Bambu package, Bambu-Clump enables efficient processing of highly multiplexed samples, including unique molecular identification (UMI) deduplication, barcode handling, chimeric read correction, and compressed output formats allowing for integration with downstream tools such as Seurat (Figure 1a-g). This is achieved through three primary mechanisms. Firstly, taking a barcoded/multiplexed bam, reference annotations and the genome (Figure 1a), Bambu-Clump generates a resource-efficient representation of reads as collective read classes while maintaining the barcode information per read (Figure 1b), enabling efficient, experiment-wide transcript discovery (Figure 1c) and read to transcript assignment (Figure 1d). Secondly, Bambu-Clump performs internal demultiplexing and produces gene counts and transcript counts based on unique reads (Figure 1e). Optionally clustering can be provided (or performed using the gene counts as part of Bambu-Pipe)(Figure 1.f) to generate cluster level (pseudobulk) transcript expression quantification (Figure 1g). Thirdly, Bambu-Clump supports an efficient data structure to account for the sparsity of transcript expression matrices across all observations. Together, Bambu-Cump provides optimised data processing and representation for highly multiplexed samples to facilitate transcript discovery and quantification simultaneously.

**Figure 1.**
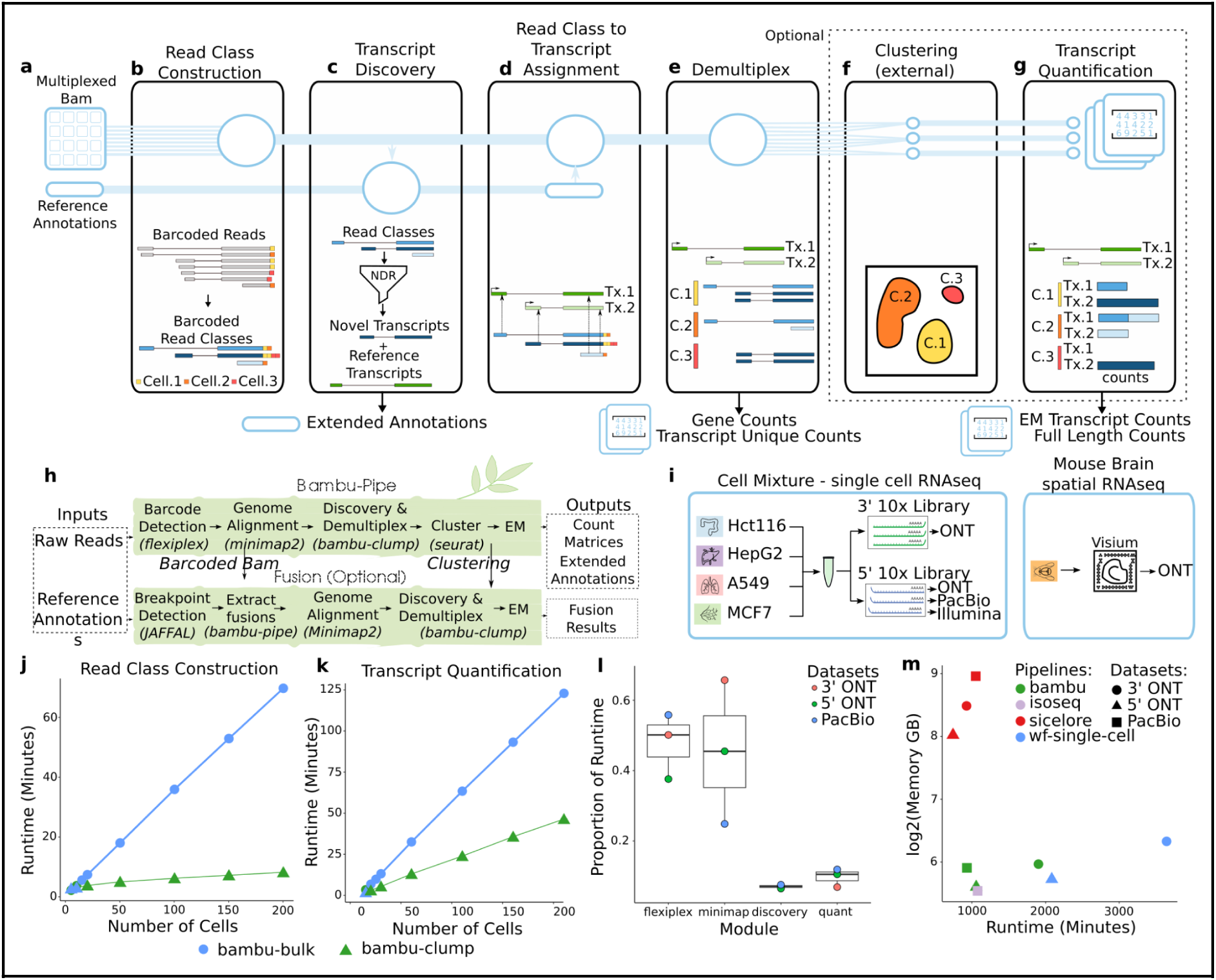
a) A graphical overview of Bambu-Clump with each step being annotated in figures b-g below. Starting from the square representing a bam file containing different barcodes, this is followed by circles representing one process, splitting into multiple based on the number of barcodes/clusters after demultiplexing. b) Read Class Construction - Bambu-Clump will load in all reads of a multiplexed bam and calculate multi-barcoded read classes. c) Transcript Discovery - Transcript discovery is done across the whole sample similar to bulk analysis, producing the extended annotations used for quantification. This process can be later repeated at the cluster level using clustering information from step f for more specialised transcript discovery. d) Read Class to Transcript Assignment - The mult-barcoded read classes are assigned to the extended annotations generating equivalence classes to be used for expectation maximisation directed quantification. e) Demultiplex - Each equivalence class is separated by barcodes producing barcode-level gene counts and unique read counts. f) Clustering - Barcodes are clustered based on gene counts with provided resolution. Clustering can also be provided for alternative methods. The default clusters are output. g) Transcript Quantification - If provided, quantification is performed using all equivalence class counts for each cluster (pseudo bulk), or for each barcode if not provided. This produces EM transcript counts and full-length counts. h) Bambu-Clump is part of the Bambu-Pipe pipeline for raw-data to count matrix processing of single-cell and spatial data. Bambu-Pipe is comprised of four modules, Flexiplex for barcode identification, Minimap2 for genome alignment and Bambu for both transcript discovery and quantification. Below shows how the Bambu-Pipe-Fusion workflow runs in parallel to the Bambu-Pipe pipeline. Both methods take the reads as a fastq input and the fusion pipeline combines the genomic alignment from Bambu-Pipe, as well as the fusion breakpoints to generate the spanning reads and construct fusion scaffolds. The spanning reads are then realigned to the fusion scaffolds where fusion discovery, and quantification is performed. Gene-count based clustering from the Bambu-Pipe pipeline can be used to generate pseudobulk counts for the fusion genes. i) An overview of the data used in this manuscript. Four human cancer cell lines (A549, HepG2, HCT116 and MCF7) were combined (“cell-mixture sample”) and 3’ and 5’ 10x libraries were created. Both libraries were sequenced with Nanopore RNA-seq, with the 5’ 10x library also being sequenced with PacBio and Illumina short-read sequencing. A spatial library preparation was performed on mouse brain using 10x Visium and sequenced with ONT. j) Scatterplots showing the runtime needed to perform read class construction against the number of input cells from the cell-mixture sample data with 1 core for Bambu-Bulk (Red) and Bambu-Clump (Blue). Each cell consists of ∼8000 reads. k) Scatterplots showing the runtime needed to perform transcript quantification against the number of input cells from the cell-mixture sample data with 1 core for Bambu-Bulk (Red) and Bambu-Clump (Blue). Each cell consists of ∼8000 reads. l) Box plots showing the proportion of running time for each of the four modules in Bambu-Pipe across the three cell-mixture long read single-cell datasets. Points are coloured by dataset, red representing 3’ ONT, green representing 5’ONT, blue representing 5’ PacBio. m) The running time and memory requirements for different single cell pipelines, Bambu-Pipe (red), Isoseq(green), Sicelore (blue) and wf-single-cell (purple) on the three cell-mixture long read single-cell datasets, 3’ ONT (circle), 5’ ONT(triangle) and 5’ PacBio (square). These were all run with 16 cores and unlimited RAM.

To facilitate systematic processing of long read single cell data, we implemented a nextflow pipeline (Bambu-Pipe), enabling the generation of count matrices directly from raw FASTQ outputs provided by sequencing services. This pipeline consists of four modules: first, barcodes and UMIs are detected using Flexiplex ^23^ for later demultiplexing; second, the labelled reads are aligned to the genome with Minimap2 ^24^; third, Bambu-Clump is executed for read class construction, transcript discovery, and single cell quantification. Finally Bambu-Clump’s expectation-maximisation based quantification is performed either at the pseudobulk or single-cell level (Figure 1h). The Bambu-Pipe pipeline includes an optional module using JAFFAL ^22^ for detecting fusion breakpoints, allowing Bambu-Clump to identify and quantify novel fusion transcripts at the single-cell level (Figure 1h). The pipeline can also be used directly starting with demultiplexed BAM files generated by other upstream tools, offering modular compatibility to use Bambu-Clump transcript discovery and quantification with various demultiplexers and aligners.

### Resource efficient processing of highly multiplexed long read RNA-Seq samples

To systematically evaluate Bambu-Clump and Bambu-Pipe, we generated five single-cell and spatial RNA-Seq data sets, including three 10x 5’ libraries for a mixture of four cancer cell lines (“cell-mixture sample”) with distinct transcriptional profiles (A549, HepG2, HCT116, MCF7) using ONT, PacBio long read RNA-Seq and Illumina short read RNA-Seq separately, a 10x 3’ library for the same cell-mixture sample using ONT, and one 10x Visium slide for wild-type mouse brain using ONT long read RNA-Seq (Figure 1i, Supplementary Text 1, Supplementary Table 1).

To assess the performance improvement of Bambu-Clump compared to the bulk RNA-Seq version of Bambu that is optimised for high depth samples such as bulk data, we compared the run time for the read class construction (Figure 1b) and quantification steps (Figure 1d,e,g) on individual cells from the 5’ ONT dataset. Here, Bambu-Clump is run on the multiplexed bam file, while the bam file was demultiplexed into separate files for each single cell in order to run the bulk version of Bambu. Bambu-Clump was consistently faster for both read class construction and quantification, reducing the average run time for 200 cells by 90% and 60% respectively, while also avoiding the need to store hundreds of individual bam files (Figure 1j,k). The runtime improvement was more pronounced with an increasing number of cells, illustrating the efficiency improvements of Bambu-Clump (Figure 1j,k). Among the four modules of Bambu-Pipe, we found that the barcode detection, alignment, transcript discovery, and quantification modules comprised 47.8%, 45.4%, 7%, and 9.7% of the runtime respectively, across the three long-read cell-mixture datasets (Figure 1l).

Next, we benchmarked the running time and memory usage of Bambu-Pipe against existing pipelines and workflows for long-read single-cell RNA-Seq, including Sicelore ^20^, ONT’s wf-single-cell ^15^, and PacBio’s IsoSeq ^18^, using the cell-mixture datasets. Generally the pipelines are either optimised for running time, or memory usage. Sicelore achieved the fastest running time (743 mins), with bambu-pipe taking (932 mins). However, Bambu-Pipe required significantly less memory across the datasets (57.2 GB for Bambu-Pipe vs 373.3GB for Sicelore) (Figure 1m). Together, these results show that Bambu-Pipe provides a resource efficient workflow for highly multiplexed long read RNA-Seq samples.

### Bambu-Clump accurately identifies new transcripts in single-cell data and is comparable to short-read methods for gene quantification

A key feature of long read RNA-Seq is the ability to discover missing transcripts, which has also been shown to improve quantification^13^. To evaluate the performance of the different methods, we randomly removed 50% of the transcript annotations from chromosome 1 of the reference GTF. We then performed transcript discovery and estimated precision and sensitivity on the removed transcripts. Only Bambu-Clump, wf-single-cell, Isosceles, and Isoseq completed transcript discovery, with Isoseq only applicable to the PacBio data (Supplementary table 2). When default parameters were used, Bambu-Clump had the highest F-score of (0.061%) compared to Isoseq and wf-single-cell with (0.015 and 0.055 respectively) (Figure 2a). However, while default parameters for Bambu-Clump prioritize precision, default parameters for wf-single-cell and Isoseq prioritize sensitivity, highlighting that default parameters may not align with research needs that may need to either prioritize sensitivity or precision (Figure 2b,c). Unlike wf-single-cell and Isoseq which do not provide options to modulate transcript discovery precision and sensitivity, Bambu-Pipe uses the novel discovery rate (NDR) that provides a continuous threshold to adjust the sensitivity and precision of transcript discovery depending on the experiment’s needs. By adjusting the NDR parameter Bambu-Pipe achieved the same level of sensitivity as the other pipelines, with greater precision (Figure 2d). Bambu is also able to target specific barcodes and clusters to improve transcript discovery of cell-type specific transcripts.

**Figure 2.**
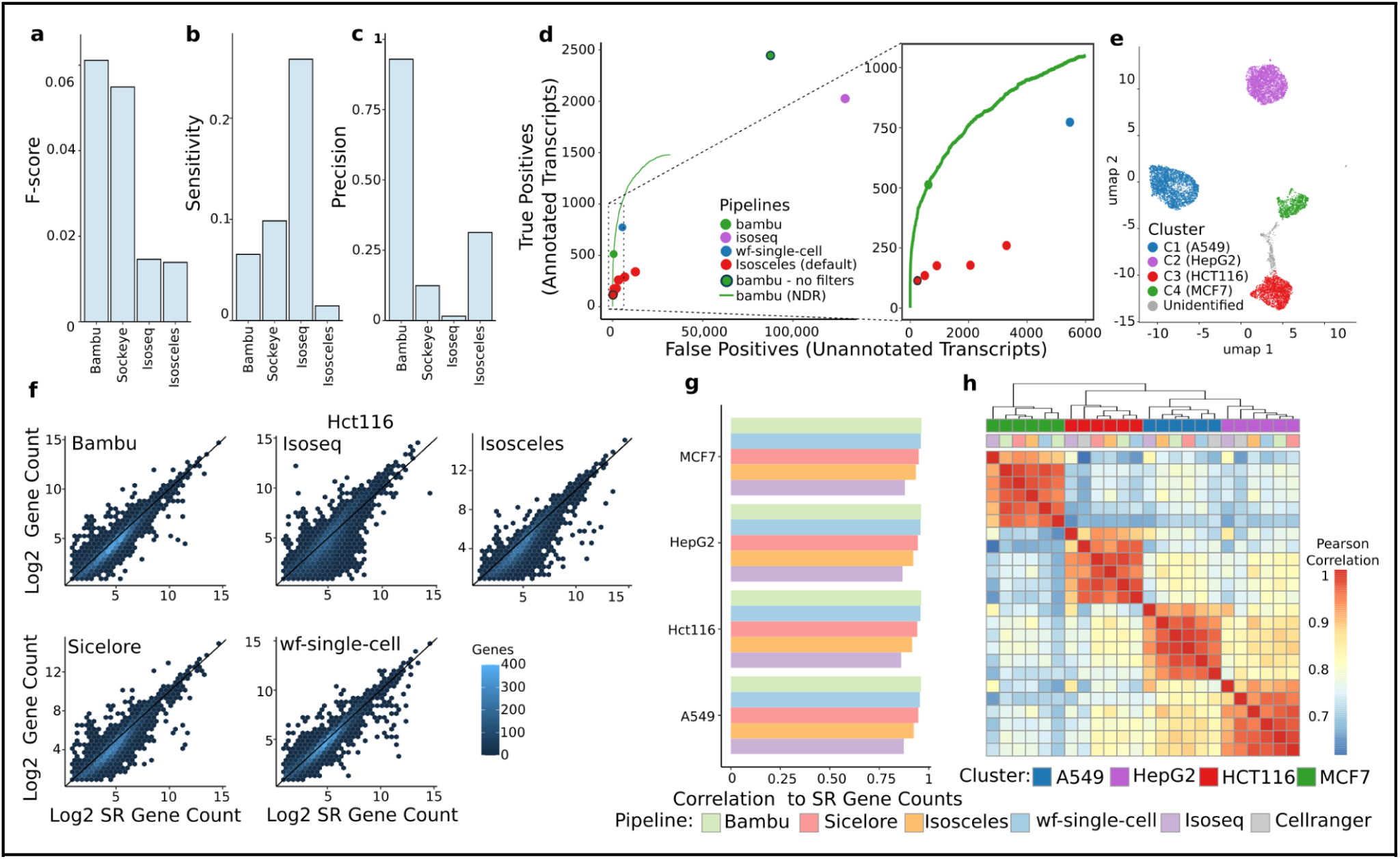
Benchmarking Bambu-Clump. **a-c)** Bar plots showing the F-score (a), sensitivity (b) and precision (c) for transcript discovery with Bambu, wf-single-cell, Isoseq, and Isosceles calculated on the 50% transcripts sampled from chromosome 1 of the 5’ Pacbio dataset. This is the default parameters for Bambu and Isosceles. **d)** A scatter plot showing the number of true positive transcripts (annotated transcripts sampled before analysis) that were discovered vs the number of unannotated transcripts that were reported of the 5’ Pacbio dataset. Bambu is shown in blue, and the line represents the different results as the NDR threshold is adjusted. A black outlined point represents Bambu run with the default additional filters turned off, the NDR threshold is 1, and the minimum gene proportion threshold is set to 0 from the default of (<0.5) and subset transcripts are allowed. Isosceles is shown in red, with its default output outlined in black (read count threshold = 1, gene proportion threshold = 0). Additional red points represent additional read count (2,5,10) and gene proportion thresholds (0.025, 0.05). wf-single-cell is represented in green and isoseq in purple. The left plot is an inset of the right plot showing a zoomed in view of the region highlighted by the dotted box. **e)** UMAP plot showing the clustering of barcodes using gene counts from bambu on the 5’ Pacbio dataset. The points are coloured based on their closest correlating cellline from bulk data, blue, purple, red and green are A549, HepG2, Hct116 and MCF7 respectively. Barcodes that fell outside of these clusters, are detected as doublets and were labelled as unidentified. **f)** Density hex plots showing the log_2_ transformed gene counts of the five pipelines against the gene counts from Cellranger of the Hct116 clusters, from left to right, Bambu, Isoseq, Sicelore, wf-single-cell, and Isosceles. The colour of the hexgon represents the number of genes whose value falls in that area shown in the legend (bottom right). g) A barplot showing the pearson correlation values of the pipeline derived gene counts against Cellranger gene counts for each cluster. The pipelines used are represented with pastel blue, orange, purple, red and green representing Bambu, Isosceles, Isoseq, Sicelore and wf-single-cell respectively h) A heatmap showing the pearson correlation values of the gene count correlation between the pipelines and clusters. Clusters are annotated in the top annotation row with blue, red, purple and green representing A549, Hct116, HepG2, and MCF7. The pipeline used is represented in the bottom annotation row with pastel blue, orange, purple, red, green, and grey, representing Bambu, Isosceles, Isoseq, Sicelore, wf-single-cell, and Cellranger respectively.

Using the single-cell generated gene counts from each tool we clustered the cells, noting similar barcode groupings for each pipeline (Figure 2e, Supplementary Text 2, Supplementary Figure 1). For tools without demultiplexing we used barcode cluster assignments from Bambu-Pipe. From these clusters we generated pseudo-bulk gene expression estimates by summing together the counts from each cell. First, we compared the gene expression estimates from the long read methods to the matching gene expression estimates from the short read RNA-Seq data. Across the four cell lines, all methods achieved a high correlation (average pearson correlation coefficient > 0.868), with Bambu showing the highest correlation to the Cell Ranger estimates (*ρ=*0.952) (Figure 2fg). Pseudo-bulk gene expression estimates from Bambu, Sicelore and wf-single-cell were most similar to each other, with Isoseq being most distinct (Figure 2h). Differences between cell types were larger than differences between methods and sequencing technologies, indicating a high level of consistency in analysing single cell gene expression data (Figure 2h). In contrast, gene expression estimates showed larger differences when compared to bulk RNA-Seq data. This observation was also seen when using the other long-read single-cell methods, and short-read single cell data (Supplementary Figure 2), suggesting that this could be related to differences in the experiment, cell lines, or library preparation methods.

### Single-cell isoform level quantification with Bambu-Pipe

Long-reads enable the quantification of single-cell data at the isoform level, which is not possible with short-read methods. Therefore, to obtain a benchmark data set for evaluating single cell transcript quantification, we simulated a single-cell experiment derived from bulk RNA-Seq data for the 4 cell lines. We generated a list of barcodes from the 10x single-cell data, and randomly assigned each of them to one of the four bulk datasets. Reads from each bulk dataset were then assigned barcodes from their allocated set, so that the mean and standard deviation of barcode counts in the simulated data matched the 10x single-cell data (Figure 3a). First we compared gene expression estimates from the simulated single-cell experiment to the ground-truth estimates obtained from the original bulk samples. We found that gene expression estimates showed a high correlation, with Bambu-Clump achieving an average Pearson correlation score of 0.93 across the four cell-lines with the majority of methods performing similarly (Figure 3b), revealing that all methods can accurately capture gene expression.

**Figure 3.**
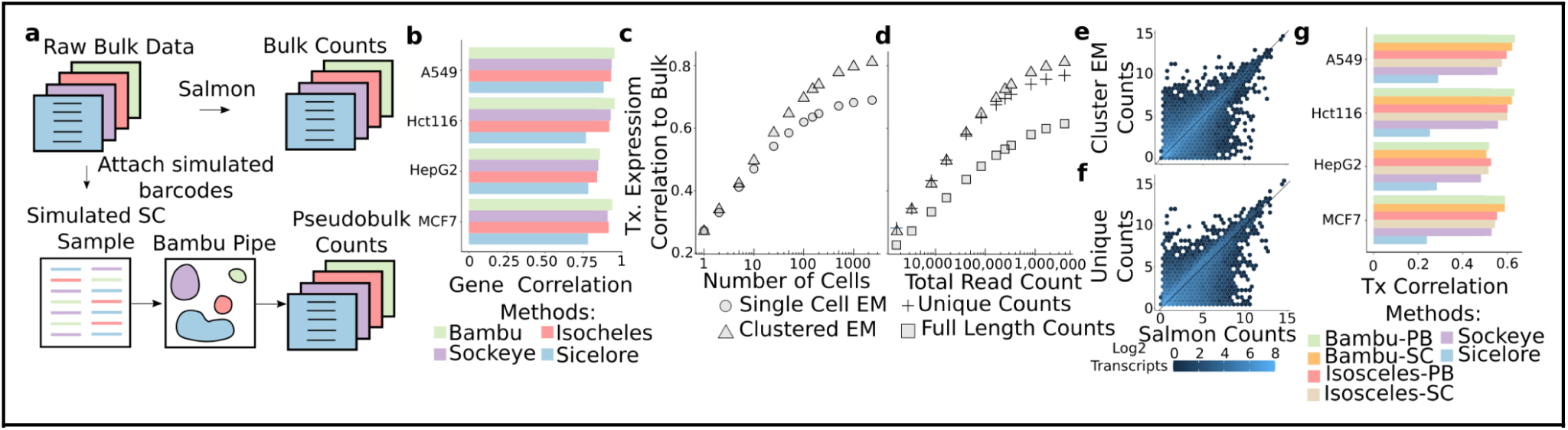
Simulating transcript data to optimise pseudobulk and single-cell level transcript quantification. **a)** To generate and evaluate simulated data, we attached barcodes to four bulk datasets and combined them together to represent four clusters. Using these clusters we compare the pseudobulk counts to the counts derived from the original individual bulk datasets. **b)** A barplot showing the pearson correlation values of the pipeline derived gene counts against Salmon gene counts run on the bulk data for each cluster. The pipelines used are represented with pastel blue, purple, red and green representing Bambu, wf-single-cell, Isosceles, and Sicelore respectively **c, d)** Scatter plots showing the pearson correlation of transcript expression for a sample of cell (c) and reads (d) from the A549 simulated cluster. The count metrics used are the EM counts from each cell (circle), performing the EM using all reads from the sample (triangle), unique counts (plus symbol), and the full-length counts (square). **e,f)** Density hex plots showing the log_2_ transformed cluster EM transcript counts (top) and unique transcript counts (bottom) of the full simulated A549 cluster against the transcript counts from Salmon run on the A549 bulk data. The colour of the hexagon represents the number of genes whose value falls in that area shown in the legend (bottom right). **g)** A bar plot showing the spearman correlation values of the pipeline derived transcript counts against Salmon transcript counts run on the bulk data for each cluster. The pipelines used are represented with pastel blue/purple, red/green, orange, and brown, representing Bambu, wf-single-cell, Isosceles, and Sicelore respectively. PB refers to transcript counts generated at the cluster level (pseudobulk) and SC represents the sum of single cell counts respectively.

Transcript quantification introduces the challenge of ambiguous assigning reads, which can be assigned using an expectation-maximisation (EM) algorithm. However, one of the key differences between bulk and single-cell quantification is the lower amount of reads per transcript per cell. To address this, Bambu-Clump includes the option to estimate transcript expression after clustering, thereby increasing the data available to the EM. To evaluate the impact of low read counts on transcript quantification, we compared transcript expression estimates from simulated single cell data at different cell quantities with the expected expression levels. We first estimated expression with EM for each single cell, and then summarised the counts for each cluster, and compared that against clustered EM. The clustered EM always performs better than the single cell EM, with the difference growing with the number of cells in the cluster (Figure 3c). The clustered EM showed good performance compared to the ground truth, being more similar to transcript quantification estimates from bulk data compared to Isosceles, wf-single-cell and Sicelore (Figure 3e,g)

It is not always possible to obtain sufficiently sized clusters to accurately perform clustered EM quantification such as from rare cell types, or transitioning cells. Therefore, we evaluated which of the single-cell level counts produced by Bambu-Clump (full-length, unique, single cell EM counts) are most accurate in low read depth situations. We found that when quantifying cells with less than ∼42k reads, unique counts were faster to calculate and more similar to the ground truth than the single cell EM counts, however cells with higher read counts were more accurately quantified with the EM (Figure 3d). We find that using only full-length counts had the lowest correlation at all tested sequencing depths, reflecting that non-full length reads are highly informative for transcript quantification (Figure 3d). When unique counts are used to quantify expression of cell clusters, we observed a minor drop in performance compared to the clustered EM, and they also out-performed other tools that also use single-cell based quantification (Figure 3f,g). While the clustered EM shows overall the highest accuracy, these results show that unique read counts provide a fast and reliable estimate for transcript expression in single cells.

### Comparing different long-read single-cell library preparation methods and sequencing technologies for transcript discovery and quantification

Using Bambu-Clump, we compared differences in sequence technologies and library preparation methods. We find that the 5’ samples have more full length reads, with PacBio having lower barcode errors (Supplementary Text 3, Supplementary Figure 3). Next we compared the performance of transcript discovery across the samples where we observed that both 5’ samples outperformed the 3’ ONT sample, with the 5’ PacBio sample showing the highest sensitivity at comparable levels of precision (Figure 4a). Despite similar levels of performance, we observed large differences in the ranking of transcripts between ONT and PacBio 5’ samples (Figure 4b). Therefore, while 5’ enrichment has an impact on novel transcript precision, the sequencing technology impacts the identity of the transcripts that are identified.

**Figure 4.**
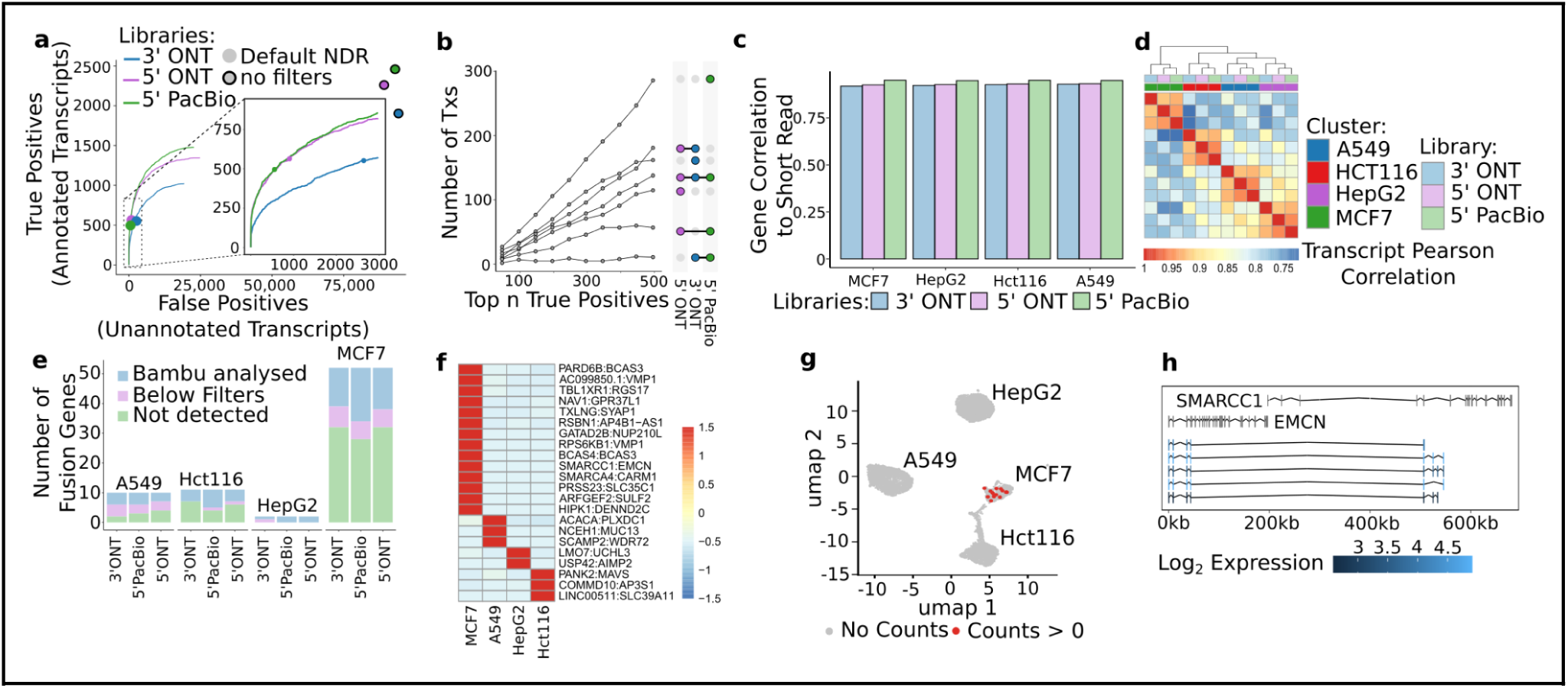
Contrasting library preparation and sequencing technology. **a)** A scatter and line plot showing the number of true positive transcripts (annotated transcripts sampled before analysis) that were discovered vs the number of unannotated transcripts that were reported from each of the samples. The line represents the different results as the NDR threshold is adjusted. A black outlined point represents when the NDR threshold is 1, the minimum gene proportion threshold is set to 0 from the default of (<0.5) ,and subset transcripts are allowed. The 3’ ONT, 5’ ONT and 5’ PacBio data are represented in blue, purple and green respectively. The left plot is an inset of the right plot showing a zoomed in view of the region highlighted by the dotted box. **b)** A scatter upset plot showing the number of true positives uniquely found in the permutation of samples when considering the top true positives. The permutations are found in the legend on the right, where a coloured circle indicates this sample is included in the grouping. The permutations are associated with the line it is directly next to. **c)** A bar plot showing the pearson correlation values of the pipeline derived gene counts against Cellranger gene counts for each cluster. The 3’ ONT, 5’ ONT and 5’ PacBio data are represented in blue, purple and green respectively. **d)** A heatmap showing the pearson correlation values of the gene count correlation between the pipelines and clusters. Clusters are annotated in the bottom annotation row with blue, red, purple and green representing A549, Hct116, HepG2, and MCF7. The samples used are represented in the top annotation row with pastel blue, purple, green, representing 3’ ONT, 5’ ONT and 5’ PacBio data respectively. **e)** Barplots showing the number of previously detected fusion genes that were able to be analysed by Bambu (blue), found with JAFFAL, but not suitable for Bambu (purple), or would not be found in the sample at all (green). This is separated by the sample and the cell-line the fusion was identified in. **f)** A heatmap of the gene expression of the fusion genes identified in the 5’ONT sample. Cells are shaded based on the z-score gene expression scaled by gene. **g)** The umap from Figure 2e where barcodes/cells are coloured red if it contained > 0 counts for the *SMARCC1*:*EMCN* fusion gene. Cells were coloured grey if they contained no counts for this fusion gene. h) The gene models (top) of *SMARCC1* and *EMCN* on an artificially constructed fusion scaffold used for fusion analysis, with the fusion transcripts (bottom) from *SMARCC1*:*EMCN*. The fusion transcripts are coloured based on their log_2_ expression in MCF7 cells.

Next we correlated the Bambu-Clump gene expression estimates from each library against the CellRanger short-read data. The 5’ libraries correlated higher than the 3’ data, with the PacBio library achieving the highest correlation with short read data across all 4 cell lines (Figure 4c). Similarly the 5’ samples were more similar to each other in their transcript quantification, than the 3’ library, suggesting that end enrichment impacts transcript-level quantification (Figure 4d).

### Fusion transcript discovery and quantification in single cells

A key advantage of long read RNA-Seq is the ability to sequence complete transcripts, enabling the discovery of fusion transcripts from single cell data. Here, we applied the fusion mode of Bambu-Pipe to the cell mixture data and compared the results to previously reported and validated fusion genes ^11^. All three libraries were able to detect similar numbers of fusions, with the 5’ PacBio library being able to quantify the highest number of validated fusions (39.7%) (Figure 4e). Individually each sample reported an additional number of candidate fusions that are likely due to randomly chimeric reads and which were largely removed when the intersection from all protocols was analysed, resulting in 74 fusions out of which 29 have previously been reported. Both the fusion genes that were undetected, and those which were not previously reported had lower read support in the original data and single cell data respectively (Supplementary Fig 4). The identified fusions were specific to their cell-line of origin suggesting that the multiplexing of the fusion samples does not impair this analysis (Figure 4f). One such example is the *SMARCC1*:*EMCN* fusion which was previously detected in MCF7 and was only found in cells from the MCF7 cluster (Figure 4g,h). We identified 5 different isoforms which consist of four exons from the *SMARCC1* gene, 1 exon from *EMCN*, and then a combination of 3 uncataloged exons from within the intron region of *EMCN*, highlighting the power of long read RNA-Seq to identify fusion transcripts from single cell RNA-Seq data (Figure 4g-h).

### Using Bambu-Clump to detect unannotated marker isoforms in spatial data

To illustrate how Bambu-Clump can be used to analyse spatial long read RNA-Seq data, we analysed 10X Visium data from a mouse brain (Figure 5a). By comparing the spatial long read RNA-Seq data to a reference data set ^25,26^, we identified 5 distinct brain regions annotated as Pallium-Glut, NN-IMN-GC, Subpallium-GABA, TH-EPI-Glut, and HY-EA-Glut-GABA (Figure 5b, c). We performed differential expression analysis and identified the top 75 putative cell marker transcripts for each region using transcript-level counts (Figure 5d). We compared our cell markers to a reference panel of 500 brain variable transcripts ^25,26^, finding an agreement of 59 transcripts with an additional 5 being transcripts from panel genes. Furthermore, we identified five novel transcripts that act as cell markers in our sample (Figure 5e). Even with long-read RNA sequencing, RNA degradation or sequencing artefacts can result in the identification of isoforms that are purely supported by fragmented reads. To enable the identification of genes that are most reliably quantified, Bambu-Clump provides unique and full-length read counts per transcript ^13^. Here, we find that 45% of marker isoforms are supported by full-length reads, providing a way to prioritise candidate isoforms for downstream analysis and experiments (Figure 5f).

**Figure 5.**
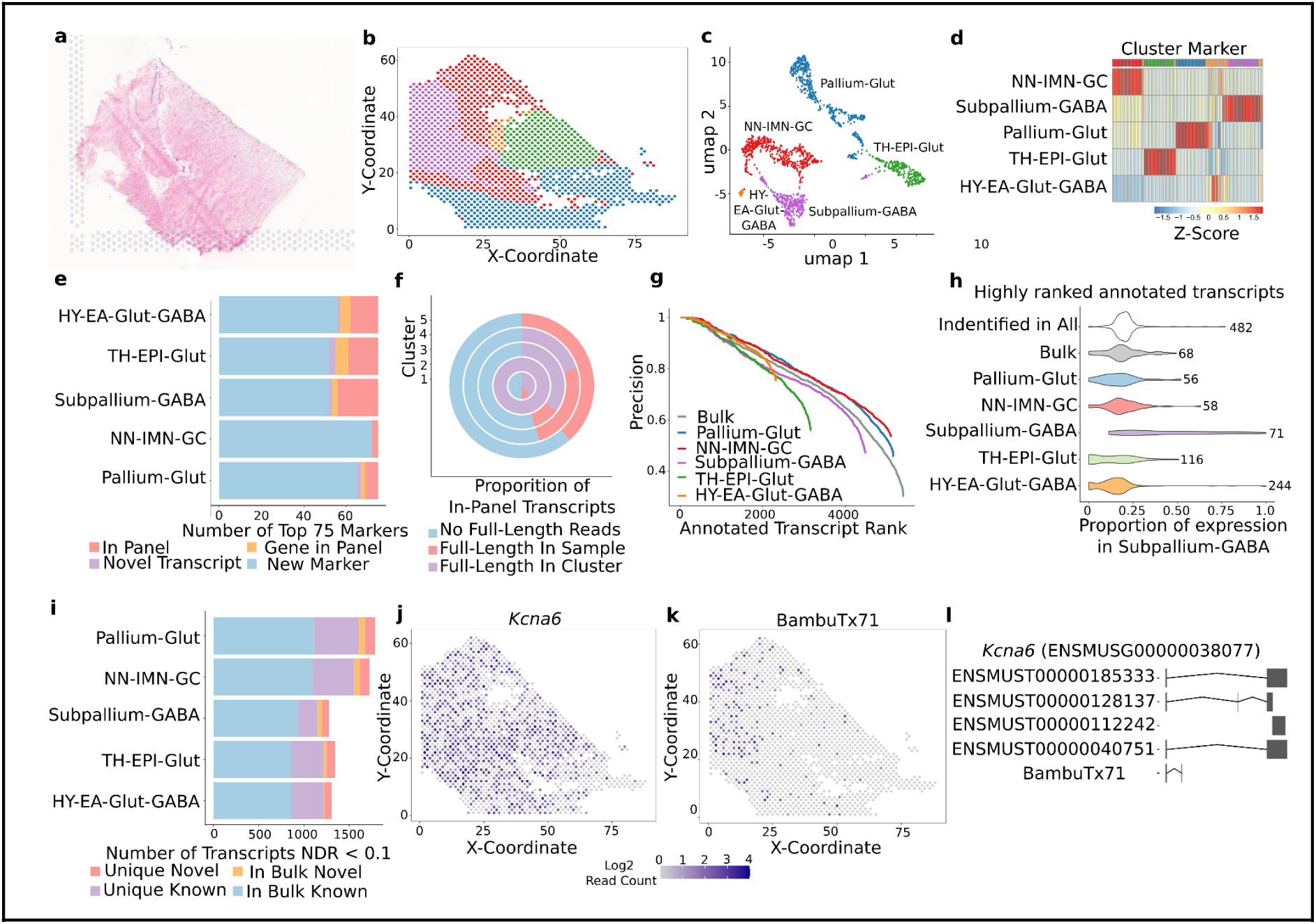
Applications of Bambu-Pipe for spatial analysis **a)** A digital photo of the mouse brain sample on the 10x slide **b)** A scatterplot showing the detected barcodes at their X and Y coordinates. Each point is coloured based on their assigned cluster based on gene counts. **c)** A UMAP of the spatial barcodes. Each cluster was annotated from the predominant annotation of each barcode in the cluster. Pallium-Glut, NN-IMN-GC, Subpallium-GABA, TH-EPI-Glut and HY-EA-Glut-GABA are coloured in blue, red,purple, green, and orange respectively. **d)** A heatmap of the top 20 marker transcripts identified for each of the 5 clusters. The colour represents the z-score which was normalised by transcript. **e)** A bar plot showing the number of marker transcripts within the top 75 from each cluster that were present in the brain atlas panel (red), was part of a gene in the panel (orange), a novel transcript detected by bambu (purple), or a marker not present in the panel (blue). **f)** A pie chart representing the proportion of marker genes found in the panel (e, red) that had full-length read support in the same cluster (purple), at least full-length read support in the whole sample (red) or had no full-length read support at all (blue). Read support is defined as having at least one full length read assigned to the transcript. **g)** A line plot showing at each rank of true positive transcripts classified during transcript discovery, what proportion of the candidate transcripts are true positives (precision). This was performed separately on each cluster with Pallium-Glut, NN-IMN-GC, Subpallium-GABA, TH-EPI-Glut and HY-EA-Glut-GABA being coloured in blue, red,purple, green, and orange respectively. The original ranking, done on all data together (bulk) is in grey. **h)** A violin plot of the proportion of transcript expression for each transcript is found in the subpallium-gaba cluster within the sample. Each violin plot represents the transcripts that were uniquely ranked in the top 1000 during transcript discovery on each cluster, or in bulk. ‘All’ represents the transcripts uniquely found in all clusters and bulk together. The number represents how many uniquely ranked transcripts comprise each violin plot. i) A bar plot showing the number transcripts achieving an NDR score below the threshold for each cluster. Transcripts that are already known and were also found in the bulk are coloured blue, the novel transcripts that were also discovered in bulk is coloured in orange. Transcripts that were below the NDR but not detected in the bulk and were unique to the cluster specific discovery are coloured in purple for known transcripts, and red for novel transcripts. **j-k)** A scatterplot showing the detected barcodes at their X and Y coordinates. Each point is coloured based on the log2 expression of ENSMUSG00000038077 (j) or a novel transcript from this gene BambuTx71 (k). **l)** The transcript models of the transcripts belonging to ENSMUSG00000038077. The Black rectangles represent exons, with the lines connecting them representing the introns between them.

Next we performed targeted transcript discovery for each brain region separately, where we observed a higher precision for the two largest regions compared to bulk (Figure 5g). To understand which transcripts are specifically identified in the targeted discovery approach versus bulk-approaches, we compared their expression across the different brain regions. While transcripts which were highly ranked by Bambu-Clump in the bulk-level discovery approach were constitutively expressed across the sample, targeted discovery ranked region-specific expressed transcripts more highly (Figure 5h). Using this targeted approach we identified 367 novel transcripts which were exclusive to targeted discovery (Figure 5i). Combining marker identification and targeted discovery we identified a full-length read supported novel isoform from *Kcna6* (ENSMUSG00000038077), whose gene is expressed across the sample, but which is uniquely expressed in the subpallium (Figure 5j-i), highlighting the potential of clustered transcript discovery in Bambu-Clump to detect novel cell markers that would otherwise be missed.

## Discussion

With the recent availability of long-read single-cell sequencing data, several computational tools have emerged for the analysis of this complex data type ^15–20^. Bambu-Clump, in conjunction with its complementary pipeline Bambu-Pipe, aims to enhance analysis at both single-cell and cluster levels by enabling targeted transcript discovery and implementing efficient methods for pseudo-bulk and single-cell quantification. Notably, Bambu-Clump expands analytical options, offering full-length counts for marker transcript validation, spatial analysis integration, and the detection and quantification of fusion transcripts.

A major challenge in single-cell transcript quantification arises from the typical low read depths, which can be partially addressed with clustered expression analyses ^21^. However, this approach does not provide single cell transcript expression estimates. While the EM can be applied to single cells as well, we observed that such methods ^13,15,21,27^ show lower performance at typical single-cell read depths. In our simulations, unique counts performed comparably to, and sometimes better than, EM-based counts at low sequencing depths. A future approach could blend these strategies, selecting counts based not on cell read depth but on gene expression depth, with EM-based quantification applied to highly expressed genes within single cells, where the clustered EM does not provide sufficient benefits.

A remaining challenge in the development of long-read single-cell RNAseq tools is benchmarking performance. While the outputs of different methods can be compared against a reference data set such as short read or bulk RNA-Seq, the lack of ground truth expression estimates limit the ability to estimate the gene and transcript quantification accuracy. Furthermore, different methods are more suitable for different types of data, for example Isosceles performs de novo transcript analysis whereas methods such as Bambu use existing annotations to improve transcript discovery. Here, we used a mixture of sampling, orthogonal data and simulations to benchmark Bambu-Clump and other methods. However, while this comparison measures the similarity to expected quantification estimates, it still does not directly measure accuracy for single cell expression estimates. Additional single cell and spatial benchmarking data sets or community-driven benchmarking efforts ^28^ could help to address this and improve the ability to quantify transcript expression for low depth RNA-Seq data An advantage of long read RNA-Seq is the presence of full length reads, which can be used to identify complete fusion transcripts, novel isoforms, and alternative splicing events in single cell and spatial data. Here we demonstrate this in single cell RNA-Seq from cancer cell lines, where the presence of fusion transcripts can indicate cancer development and progression ^29–32^, and spatial data from mouse brain, isoform expression has been associated with neurodegenerative diseases ^33^. Together, Bambu-Clump and Bambu-Pipe present easy-to-use, efficient, and accurate transcript discovery and quantification for highly multiplexed long-read RNA-seq datasets. By supporting single cell and clustered quantification and transcript discovery, Bambu-Clump can be adapted to varying read depths, barcode complexities or the analysis of rare cell types. Combined with the ability to provide unique, full length, and EM based expression estimates, Bambu-clump provides rich information from long read RNA-seq to drive new insights into the role of transcript expression in single cell and spatial biology.

### Software and Data Availability

Bambu-Clump is available on github at https://github.com/GoekeLab/bambu/tree/Bambu-Clump_biorxiv

Bambu-Pipe is available on github at https://github.com/GoekeLab/bambu-singlecell-spatial The single-cell and spatial data is available at https://www.ebi.ac.uk/ena/browser/view/PRJEB83819

## Methods

### Cell culturing and mouse brain extraction

A549, HepG2, HCT116 and MCF7 cells were obtained from the American Type Culture Collection (ATCC) and passaged in Dulbecco’s Modified Eagle Medium (DMEM, Life Technologies) supplemented with 10% Fetal Bovine Serum (FBS, Hyclone) in the presence of penicillin/streptomycin. Cells were dissociated using 0.05% Trypsin-EDTA (Life Technologies) and counted using the Countess II Automated Cell Counter (Thermo Scientific).

18 months old mouse cerebrum were harvested and cut into 2 equal halves along the mid-sagittal plane. The left half was embedded and cryopreserved in OCT. 10 µm cryosections were placed on 10x Visium Spatial Gene Expression slides for fresh-frozen tissues and processed following the Manufacturer’s protocol (Visium Spatial Gene Expression Reagent Kits, CG000239, Rev G).

### Library Preparation and Sequencing

Unfragmented cDNA pooled from four human cancer cell lines—colon (HCT116), lung (A549), liver (HepG2), and breast (MCF7)—was sequenced using the 10X Genomics 3’ and 5’ single-cell cDNA protocols. For nanopore sequencing, libraries were prepared using the SQK-LSK114 kit and sequenced on a PromethION platform with a runtime of 72 hours. Basecalling was performed using Guppy v6.3.8. PacBio data was generated on a Sequel IIe platform using a SMRT Cell 8M flow cell and the MAS-Seq for 10X Single Cell 3’ kit (PN: 102-659-600), with a custom oligo optimized for use with the 10X 5’ library. Short-read libraries were sequenced on an Illumina NovaSeq 6000 platform, generating 150 million reads per library.

### Code changes

Programmatically Bambu-Clump achieves a fast running time with smaller memory requirements with four foundational improvements: (1) demultiplexes reads or accepts barcoded aligned reads respectively; (2) read classes are constructed experiment wide, and not at the single-cell level; (3) data structures have been optimised to support the memory requirements of high dimensional data; (4) counts are provided at the gene level, for unique counts, and at the pseudo-bulk level for expectation maximisation counts.

### Benchmarking

To benchmark the memory and runtime performance of other long-read single-cell RNAseq pipelines we ran them on the identical cluster. These runtime stats were collected from nextflow reports, and for pipelines that did not already have a nextflow pipeline we created one that consisted of the standard commands. Each run was provided 16 CPUs, as much memory as needed and a maximum of 3 days runtime per sample. To calculate maximum memory usage for pipelines with concurrent processes, the timeline was also outputted, and the memory usage at each timepoint was calculated to find the maximum over the entire run. For the 5’ libraries the 737K-august-2016 barcodes were provided. For the 3’ library the 3M-february-2018 barcodes were provided.

Below are the parameters used for each tool used during benchmarking Isosceles - For prepare_transcripts the recommended min_bam_splice_read_count and min_bam_splice_fraction of 1 and 0.01 were used respectively. For bam_to_tcc() run_mode = “de_novo_loose”, min_read_count = 1, min_relative_expression = 0. Was run twice, once with tcc_to_transcript() to get single-cell counts and once with pseudobulk_tcc where we used the clustering from Bambu’s gene counts for the cell_labels. Isoseq - Followed the Iso-seq CLI workflow for single-cell. In summary this uses lima, isoseq, minimap2 and pigeon. --design 16B-10U-10X-T --primers primers_5prime.fasta wf-single-cell -epi2me-labs/wf-single-cell versionX) --kit_name 5prime --kit_version v1 --expected_cells 8000 Sicelore - scanfastq --bcEditDistance 1 SelectValidCellBarcode MINUMI=1 ED0ED1RATIO=1 IsoformMatrix DELTA=2 MAXCLIP=150 METHOD=STRICT AMBIGUOUS_ASSIGN=false 5’ libraries additionally had scanfastq and assignumis with --fivePbc provided To benchmark transcript discovery performance we removed 50% of the annotations from chromosome 1 of hg38 and reran all the pipelines and libraries. The 50% of annotations that were removed were considered true positives, and all other reported unannotated transcripts are considered false positives, for the purpose of this evaluation. Clustering was performed by Seurat ^34^, with a resolution of 0.05, using 15 principal components, a nFeature_RNA_threshold of 1000, and an nFeature_RNA_threshold_max of 9000. Gene-level quantification evaluation was compared against either short-read counts generated by CellRanger using the Pearson correlation for all expressed genes found in both the short-read and long-read samples. This was repeated at the transcript level using the simulated data described below.

### Simulated data

We retrieved the list of used barcodes from the 5’ PacBio 10x single-cell data. New barcode counts were sampled for each barcode using the sampled divided mean and standard deviation barcode counts. Barcodes were then randomly assigned to the following SG-Nex sample fastq files as representatives of the four cell lines: SGNex_A549_cDNAStranded_replicate5_run2, SGNex_Hct116_cDNA_replicate3_run3, SGNex_HepG2_cDNAStranded_replicate5_run4 and SGNex_MCF7_cDNAStranded_replicate3_run2. Barcodes are randomly assigned to reads from the sample until the barcode counts are greater or equal to the number of reads, with the last barcode being partially assigned and its remaining counts being unassigned. Each barcode was flanked by a random buffer for 40 nucleotides, flanking sequence 1 [‘CTACACGACGCTCTTCCGATCT’] on the 5’ end, and followed by on the 3’ end, a 10bp random UMI sequence and flanking sequence [‘TTTCTTATATGGG’] before being appended to the 5’ end of the read. The four modified fastq files were then merged together to create a simulated multiplexed input for each pipeline. To simulate single cells of different sequencing depth and clusters with different numbers of cells we sampled (1, 2, 5, 10, 25, 50, 100, 150, 200, 500, 1000 and all) barcodes 5 times each and used the associated reads for quantification comparison, using the mean correlation from the 5 replicates.

### Fusion-pipeline

Chromosomal rearrangements and structural variations can lead to the expression of fusion transcripts, which are often markers in cancer progression. Fusion transcripts can be detected in bulk-long read data (SG-Nex) however this excludes the ability to isolate the cells or progression stages in which these fusion transcripts are expressed. We added a fusion-mode to the single-cell Bambu pipeline which runs parallel using the same fastq input (Figure 4j). Using JAFFAL (ref) and the genome alignments from Bambu-Pipe it extracts the fusion spanning reads and runs them with Bambu’s fusion mode, detecting and quantifying isoforms across the breakpoints. As the spanning reads are unlikely to be sufficient for quality clustering, the clustering can be derived from the regular gene counts to do pseduobulk EM quantification of the fusion genes.

When --fusionMode is provided to Bambu-Pipe, three additional steps are included in the pipeline. The first runs JAFFAL identifying the fusion genes breakpoint regions in the sample from the raw fastq files. The next step uses the identified breakpoints to generate artificial fusion scaffolds, placing the fused genes next to each other on their own scaffold. Additionally a matching gtf file containing the transcript models for the respective genes is created. The reads which are aligned to this region are extracted from the bam file, aligned during the main alignment step of bambu-pipe, and these are realigned with minimap2 to the created scaffolds using -ax splice -G2200k -N 5 --sam-hit-only. In the final step the newly aligned fusion reads are run in Bambu with fusionMode = TRUE and NDR = 1 which generates the fusion gene counts and unique counts. An additional extra step can be performed which takes the clustering generated from the gene counts in the main workflow and calculates the pseudobulk EM counts for the fusion genes.

### Spatial

The library was sequenced twice and combined in silico. Adjustments to the Bambu-Pipe arguments were as follows: --spatial --chemistry 10x3v3. The visium-v1_coordinates.txt.gz were provided as the --whitelist which also contains the x and y coordinates. The reads were aligned to GRCm39 using the corresponding annotations. To annotate the barcodes, we converted the gene count matrix to h5ad format with the anndata R package and submitted it to the Brain Knowledge Platform MapMyCells ^25,26^ to the “10x Whole Mouse Brain (CCN20230722)” reference taxonomy and used the” Hierarchical Mapping” mapping algorithm. Instead of the class names, we annotated the barcodes using the associated neighbourhood for each class name. To cluster the spatial barcodes we used the same method as the single cell data, and displayed the clustering on top of the x and y coordinates that are stored in the Bambu output. A resolution of 0.065 was used to match the number of clusters produced by MapMyCells; other parameters were kept the same as previous clustering. The resolution was chosen to best match the resolution of clusters found by MapMyCells. Annotations were assigned to these clusters if at least 90% of the barcodes matched between the Seurat clustering and the annotations from MapMy Cells. For comparisons of marker genes we used the merfish gene panel used by the Brain Knowledge Platform. The short-read spatial data from the same sample was processed with SpaceRanger using default parameters, and clustered and analysed using Seurat’s spatial functions. Markers were identified using Seurat’s FindAllMarkers function with a min.pct of 0.25 and a logfc.threshold of 0.25. Additionally we applied the following filtered p_val_adj < 0.05 and avg_log2FC > 1 and selected the top 75 per cluster when sorted by avg_log2FC. .

### Targeted Transcript Discovery

To perform targeted transcript discovery we reran Bambu-Clump, providing the clustering generated from the initial analysis of the spatial data using the clusters argument. This produces different rankings of transcript discovery for each cluster, in addition to the original ranking which uses all cells. In order to not bias the evaluation of performance on the clusters, we did not remove annotations, and instead measured performance by comparing the ranking of annotated transcripts versus the ratio of novel transcripts identified. To compare the clusters and identify uniquely detected transcripts we both took the top 1000 ranked transcripts and all transcripts ranked with an NDR < 0.1 3

## Supplementary Text

### Supplementary Text 1 - Data description

Using Bambu-Clump’s comprehensive package of discovery and quantification, we applied it to compare the differences between 5’ and 3’ 10X libraries and sequencing with ONT and PacBio. The 10x Visium library preparation method differs from standard ONT and PacBio methods by enriching the 5’ or 3’ ends of transcripts. The impact of these experimental design decisions on the produced data, including potential biases, remains unclear. The long-read samples averaged 96 million reads, with the 5’ PacBio data containing the most (125 million) and the 5’ ONT data the least (57 million) (table 1). Our analysis revealed that 5’ ONT data exhibits the highest median read length (836bp), while 3’ ONT and 5’ PacBio data show similar mean read lengths (692bp and 675bp respectively) (figure 4a, table 1). The enrichment method or technology did not have a large impact on the number of detected cell barcodes, mapping percentage or the number of detected transcripts (table 1). The 5’ libraries contained higher proportions of full-length, spliced and uniquely-assignable reads (0.62, 0.79, and 0.79 respectively for the 5’ ONT library) compared to the 3’ library (0.48, 0.36, and 0.73) (figure 4b). This trend is also evident in transcript coverage, where the 5’ library samples show consistent coverage across the transcript body, where-as the 3’ library shows higher coverage at the 3’ end, with a loss of coverage towards the 5’ end (figure 4c). These differences highlight the larger impact of end enrichment than sequencing technology on the capture of full-length and degraded reads, however, how this impacts the quality of transcript discovery and quantification remains to be seen. To measure the accuracy of individual barcode assignment we limited our analysis to genes that were uniquely expressed in one of the four cell lines according to the bulk data, and measured the proportion of reads from these genes being counted in a different cluster. The short read data had the lowest occurrence of reads being assigned to different clusters than expected, followed by PacBio (Figure 4h). Both ONT libraries had increased proportions of aberrant assignment of counts suggesting the reads are more error prone during demultiplexing (Figure 4h).

### Supplementary Text 2 - Comparison of Gene level based clustering

Cell type annotation is a crucial step in single-cell analysis, where cells are aggregated through various methods, such as grouping them by similar gene expression patterns. To compare the clustering performance of each tool, we performed cell clustering with Seurat ^34^ using the cell-gene matrices generated by each pipeline(Figure 2e). The counts from each pipeline all resulted in 4 distinct clusters, corresponding to the 4 cell lines present in the dataset (Supplementary Figure 1). We then assigned each cluster to the corresponding cell line based on the highest correlation of gene expression with bulk RNA-Seq data ^11^. Using the short-read single cell RNA-seq data as ground truth, all pipelines assigned the majority of cells with the correct cell type (average 92.6%) (Supplementary Figure 1). Amongst the remaining cells, on average across the tools 4.6% were not detected, while 2.4% were assigned to a different cell type cluster (Supplementary Figure 1). The barcode composition of the clusters from the long-read methods also matched each other, having an average Jaccard index scores for Bambu, wf-single-cell, Sicelore and Isoseq being 0.96, 0.96, 0.95 and 0.92 respectively (Supplementary Figure 1). Overall these results show that long-read single-cell methods return similar cell clusters as established short-read methods.

### Supplementary Text 3 - Comparison of libraries

The mean transcript length for the rediscovered transcripts found uniquely by the 5’ ONT sample was the highest, whilst the transcripts found by both the 3’ ONT and 5’ PacBio samples, and 3’ ONT alone were shorter (supplementary figure 3). The 3’ ONT sample also uniquely ranked several true positives highly (figure 3i), and looking at these transcripts we saw that these had lower CPM values and contributed less to the overall gene counts, than transcripts detected by the 5’ methods (supplementary figure 3).Conversely, when looking at false positive and putative novel transcripts, these were mostly uniquely found in individual libraries, and it was less common that they were uniquely identified across multiple libraries (supplementary figure 3). Together this shows that while both 5’ samples were shown to be more effective for transcript discovery, sequencing technology has a large impact on the identity of reported transcripts.

## Acknowledgements

We thank Shyam Prabhakar, Kock Kian Hong and Tan Le Min for the preparation of exploratory data, and valuable discussions. funded by the Agency for Science, Technology and Research (A∗STAR), Singapore. J.G. is supported by grants from the National Medical Research Council (NMRC, OFIRG20nov-0108 and OFIRG16nov019), National Research Foundation NRF-NRF 08-2022-0004

## Author Contributions Statement

A.S, M.H.L, J.S, J.G conceived the project and designed the study. A.S, J.G designed the experiments. A.S, M.H.L, C.Y, J.G contributed to pipeline development, method development and data analysis. A.P, B.C, Y.X.S, E.Y.C, J.W, contributed to growing cell lines, RNA extraction, and single-cell data handling. H.L, O.B.L.A, T.W contributed to acquiring mouse brain samples and spatial sequencing preparation. A.S and J.G wrote the paper with contributions from all authors.

## Supplementary Figures

**Supplementary Figure 1.**
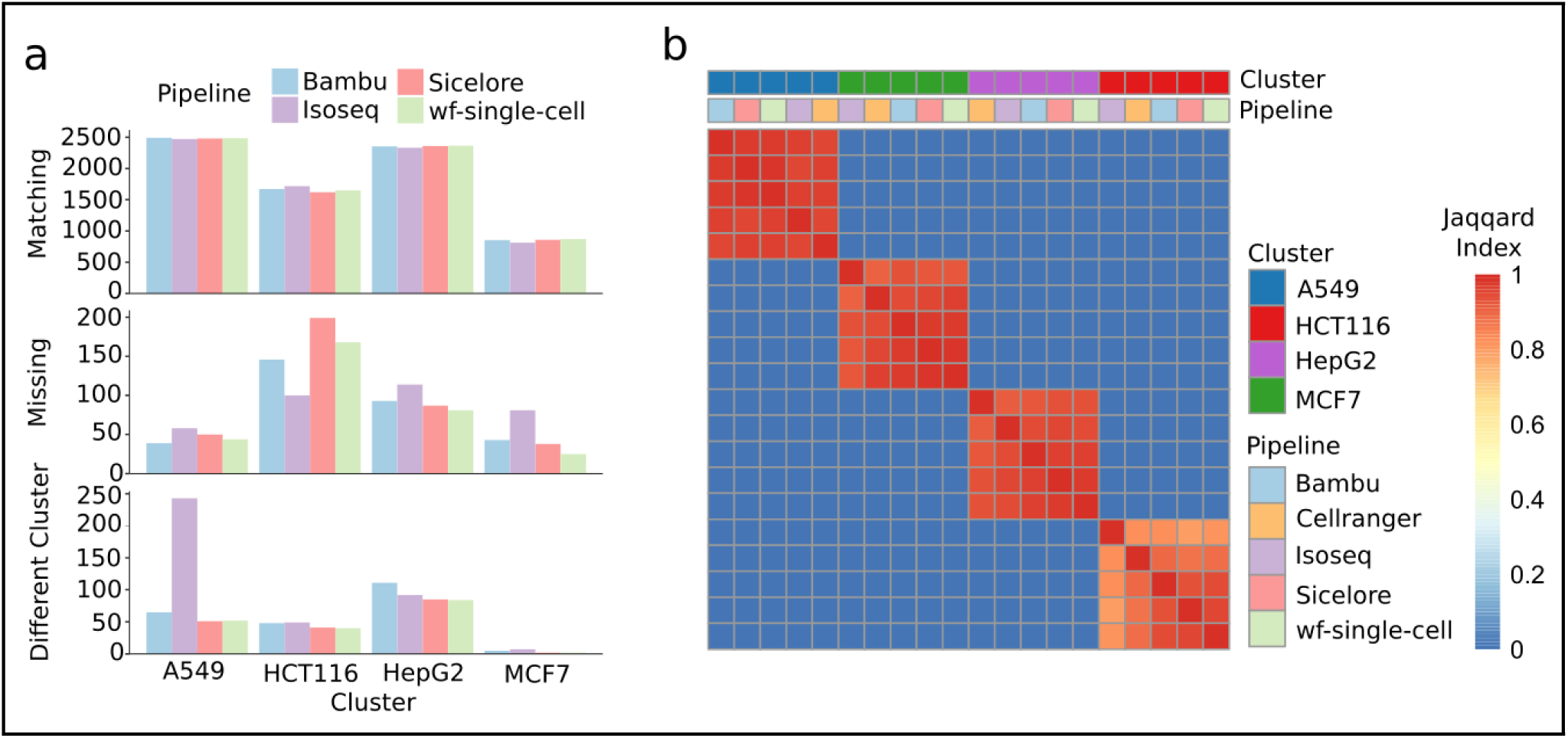
Clustering Across Tools a) Barplots comparing the barcode clustering identity between Bambu, Isoseq, Sicelore andwf-single-cell to the clustering generated using short-reads and Cellranger. The top plot shows for each method labelled in blue, purple, red and green for Bambu, Isoseq, Sicelore and wf-single-cell respectively, how many barcodes were assigned the same cluster by CellRanger. The middle plot shows the number of barcodes assigned to the cluster by CellRanger, but not by the method. The bottom plot shows the number of barcodes the methods assigned to the cluster, that CellRanger assigned differently. A heatmap showing the jacquard index of the barcode sets between each cluster and method. Clusters are annotated in the top annotation row with blue, red, purple and green representing A549, Hct116, HepG2, and MCF7. The pipeline used is represented in the bottom annotation row with pastel blue, orange, purple, red and green representing Bambu, Cellranger, Isoseq, Sicelore and wf-single-cell respectively.

**Supplementary Figure 2.**
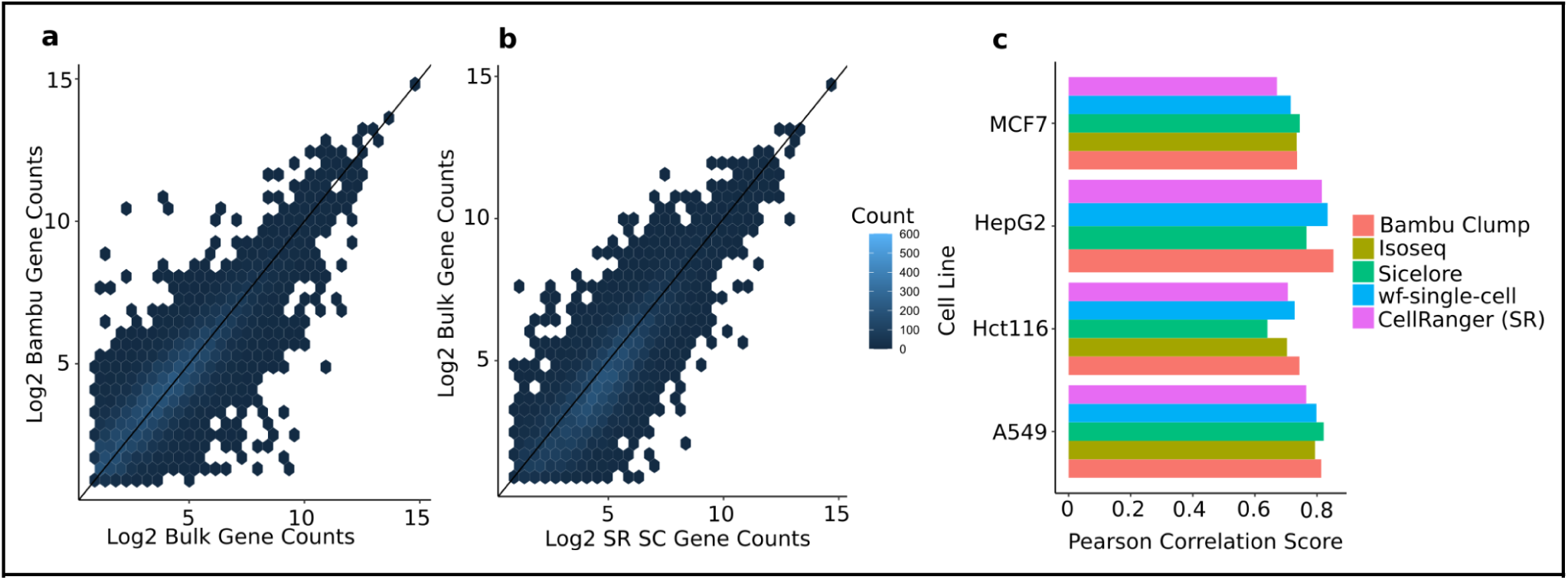
Short-Read SC vs Bulk. **a,b)** Density hex plots showing the log2 transformed Bambu-Clump gene counts vs long-read bulk (left) and short-read vs long-read bulk (right) for the A549 cluster against the gene counts from Salmon run on the A549 bulk data. The colour of the hexagon represents the number of genes whose value falls in that area shown in the legend (bottom right). **c)** Bar plots showing the pearson correlation score of gene counts from Bambu-Clump (red), Isoseq(yellow), Sicelore (green), Sockeye (Blue) and Cell Ranger on short read data (purple) against long-read bulk data quantified with Salmon.

**Supplementary Figure 3.**
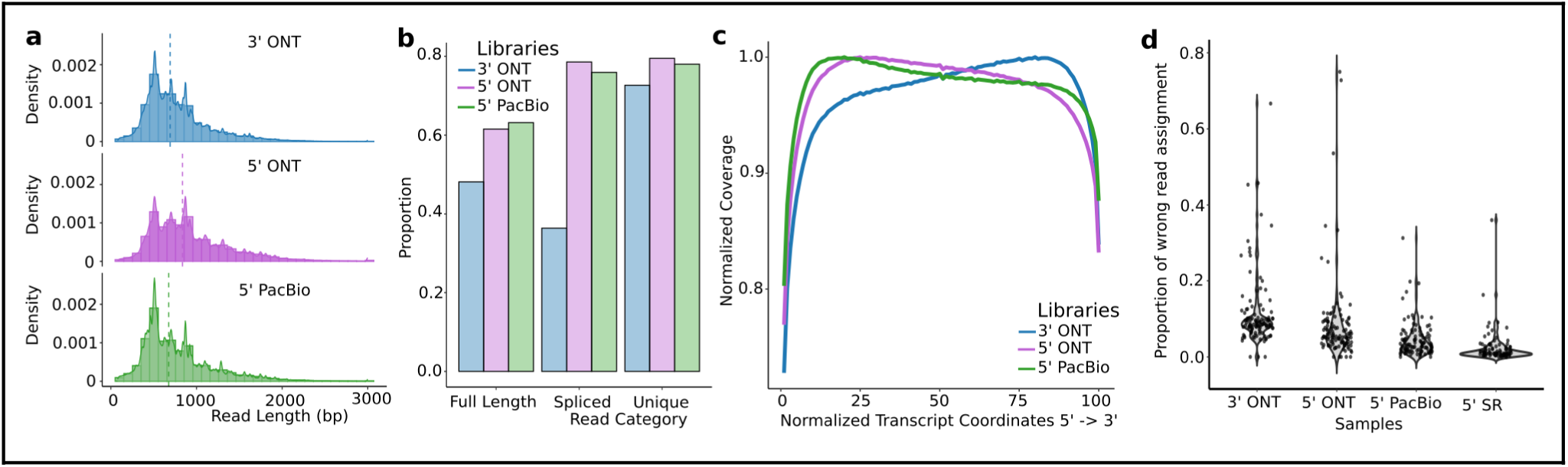
a) Histograms showing the distribution of read lengths from the three samples, 3’ ONT (top, in blue), 5’,ONT (middle, in purple) and 5’PacBio (bottom, in green). The dotted line represents the mean read length for the sample. **b)** Barplots showing the proportion of reads from each sample that are full-length (contain all splice junctions), are spliced, or can be uniquely assigned to one annotated transcript. The 3’ ONT, 5’ ONT and 5’ PacBio data are represented in blue, purple and green respectively. **c)** A line plot showing the normalised average read coverage of reads across a normalised transcript body in 5’ to 3’ direction. **d)** Violin plots of the proportion of reads from cell line specific genes being assigned to different cell lines in the single-cell data. Each point represents one gene.

**Supplementary Figure 4.**
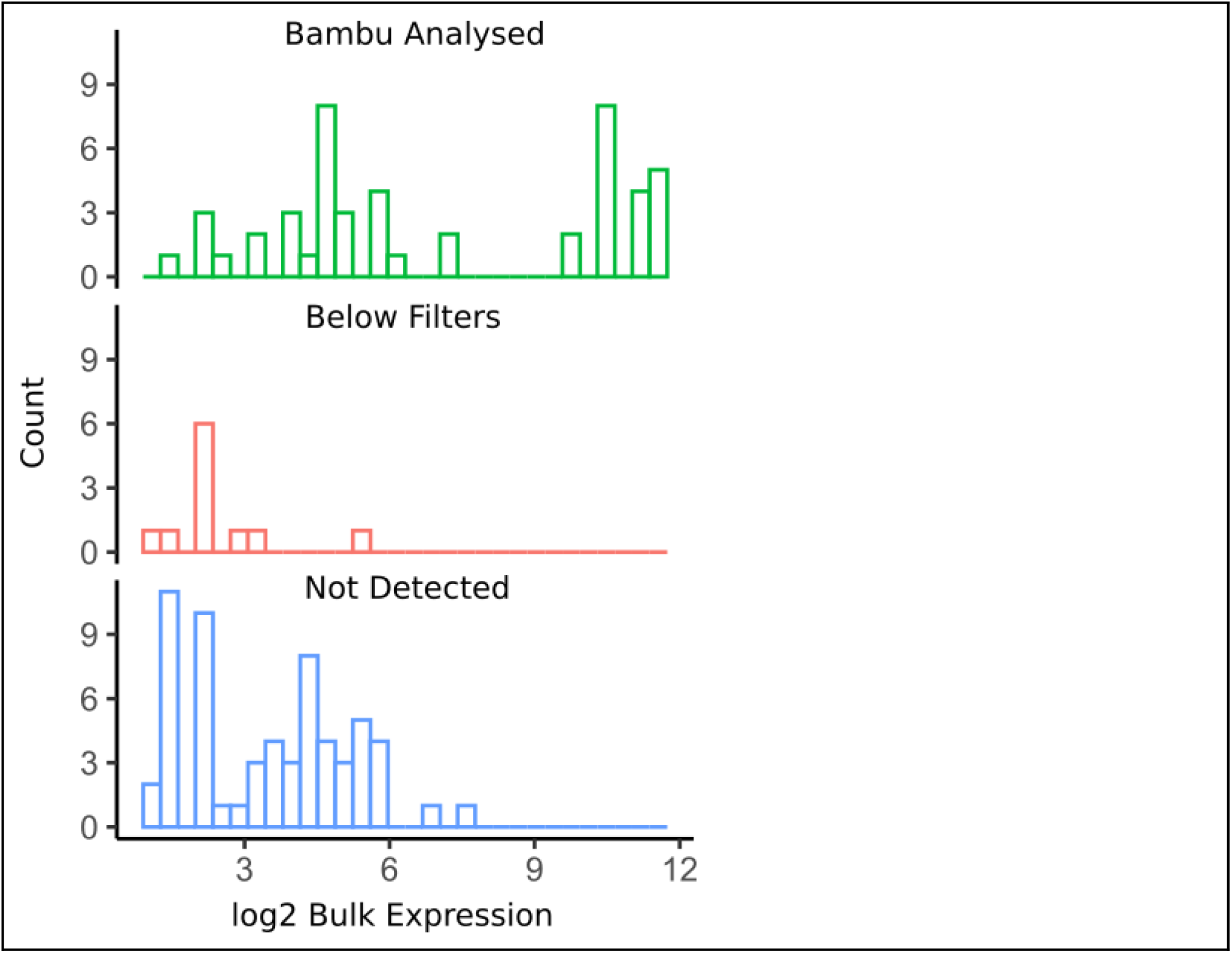
Bulk expression of detected and undetected Fusion Genes. A histogram showing the expression frequency of the fusion transcripts detected in long-read bulk and then quantified with Bambu (green, top), below Bambu’s expression filters (middle, red), or not detected by JAFFAL (bottom, blue).

---

**Supplementary Table 1.** Sample Statistics. Number of Cells is based on valid barcodes from flexiplex Transcript FL counts are Full-Splice-Match counts from Bambu

|  | 5' ONT | 3' ONT | 5' PacBio | 5' Short Read | Spatial (mouse) |
| --- | --- | --- | --- | --- | --- |
| Number of Reads | 57,551,624 | 105,064,936 | 125,617,514 | 2,714,917,074 | 108,703,288 |
| Average Read Length | 1126bp | 987bp | 852bp | 150bp (paired) | 466 |
| Mapping % | 97.1% | 95.7% | 96.1% | 88.2% | 75.4% |
| Number of Barcodes | 8933 | 8709 | 8929 | 9474 | 1512 |
| Transcripts >= 1 FL Count | 50,739 | 55,180 | 54,510 | - | 26,481 |
| Full Length Counts (FSM reads) | 32,894,248 | 44,379,912 | 66,177,002 | - | 4,562,939 |

**Supplementary Table 2.** Software used in Benchmarking.

| Tool | Version | Notes: |
| --- | --- | --- |
| Isoseq | pbbam : 2.4.99 (commit v2.4.0-16-g5cc6e4b)<br>pbcopper : 2.3.99 (commit v2.3.0-14-g5ac5693)<br>pbmm2 : 1.11.99 (commit v1.11.0-1-g1b5a417)<br>minimap2 : 2.15<br>parasail : 2.1.3<br>boost : 1.77<br>htslib : 1.17<br>zlib : 1.2.13 | Cannot run ONT |
| wf-single-cell | pysam 0.21.0<br>parasail 1.2.3<br>pandas 2.0.3<br>rapidfuzz 2.13.7<br>scikit-learn 1.3.0<br>fastcat 0.13.2<br>minimap2 2.24-r1122<br>samtools 1.17<br>bedtools v2.30.0<br>gffread 0.12.7<br>seqkit v2.5.1 | Needed to manually combine intermediate files to generate a GTF File |
|  | stringtie 2.2.1 |  |
| Sicelore | v2.1 | Requires additional reference information/Could not get to run discovery |
| Isoquant | v3.3.1 | Produces a large amount of intermediate files >5TB preventing me from running it on the HPC |
| Isosceles | Isosceles_0.2.1 | Single-cell transcript quantification for 5'PB and 3'ONT exceeded 3 day HPC runtime limit |
| Supplementary Table 2: Software used in Benchmarking |  |  |

**Supplementary Table 3.** Features of Bambu-Pipe and Bambu-Clump in the long-read single-cell ecosystem.

| Features | Bambu | IsoQuant | wf-single-cell | Iso-Seq | Sicelore | Isosceles |
| --- | --- | --- | --- | --- | --- | --- |
| Supports ONT and PacBio data | ✓ | ✓ | ✓ | ✗ | ✓ | ✓ |
| Supports long read spatial RNA-Seq | ✓ | ✗ | ✗ | ✗ | ✓ | ✗ |
| Outputs full-length counts and unique counts | ✓ | ✗ | ✗ | ✓ | ✗ | ✗ |
| Provides isoform quantification at the pseudo bulk level | ✓ | ✗ | ✗ | ✗ | ✗ | ✗ |
| Customizable transcript discovery sensitivity and precision parameters | ✓ | ✗ | ✗ | ✗ | ✓* | ✓ |
| Optimised sample specific discovery parameter | ✓ | ✗ | ✗ | ✗ | ✗ | ✗ |
| Ability to detect and quantify fusion transcripts | ✓ | ✗ | ✗ | ✗ | ✓ | ✗ |

